# PHENOS: a high-throughput and flexible tool for microorganism growth phenotyping on solid media

**DOI:** 10.1101/195784

**Authors:** David B. H. Barton, Danae Georghiou, Neelam Dave, Majed Alghamdi, Thomas A. Walsh, Edward J. Louis, Steven S. Foster

## Abstract

**BACKGROUND:** Microbial arrays, with a large number of different strains on a single plate printed with robotic precision, underpin an increasing number of genetic and genomic approaches. These include Synthetic Genetic Array analysis, high-throughput Quantitative Trait Loci (QTL) analysis and 2-hybrid techniques. Measuring the growth of individual colonies within these arrays is an essential part of many of these techniques but is useful for any work with arrays. Measurement is typically done using intermittent imagery fed into complex image analysis software, which is not especially accurate and is challenging to use effectively. We have developed a simple and fast alternative technique that uses a pinning robot and a commonplace microplate reader to continuously measure the thickness of colonies growing on solid agar, complemented by a technique for normalizing the amount of cells initially printed to each spot of the array in the first place. We have developed software to automate the process of combining multiple sets of readings, subtracting agar absorbance, and visualizing colony thickness changes in a number of informative ways.

**RESULTS:** The “PHENOS” pipeline (PHENotyping On Solid media), optimized for *Saccharomyces* yeasts, produces highly reproducible growth curves and is particularly sensitive to low-level growth. We have empirically determined a formula to estimate colony cell count from an absorbance measurement, and shown this to be comparable with estimates from measurements in liquid. We have also validated the technique by reproducing the results of an earlier QTL study done with conventional liquid phenotyping, and found PHENOS to be considerably more sensitive.

**CONCLUSIONS:** “PHENOS” is a cost effective and reliable high-throughput technique for quantifying growth of yeast arrays, and is likely to be equally very useful for a range of other types of microbial arrays. A detailed guide to the pipeline and software is provided with the installation files at https://github.com/gact/phenos.

## Background

Around the turn of the new millennium, following developments in DNA microarray technology, microbial geneticists began to work with large regularly ordered arrays of colonies of genetically distinct microorganisms. For example, *Saccharomyces cerevisiae* deletion mutant arrays were announced in 1999 [1] and made it possible to rapidly screen the whole yeast genome for synthetic lethal interactions in Synthetic Genetic Array (SGA) analyses [2, 3]. That same technique was soon expanded to other species like the fission yeast *Schizosaccharomyces pombe* [3], and the bacterium *Escherichia coli* [4], Other laboratories were also putting gene expression libraries into *E. coli* or in *S. cerevisiae* arrays for high-throughput screening [6, 7],

As these approaches gained ground, robotic colony manipulation systems became commercially viable tools to help create and duplicate these microbial arrays. For example, the VersArray^™^ Colony Picker and Arrayer Systems from BioRad Laboratories, the QPix 400 Series Microbial Colony Pickers from Molecular Devices, and the easy-to-use ROTOR HDA Colony Manipulation Robot (Singer Instruments in Somerset, UK) which was specifically conceived for SGA analysis.

In just the last few years, the ROTOR HDA system alone has been used for purposes as diverse as screening *S. cerevisiae* for thermotolerance [6], working with *S. pombe* auxotroph deletion libraries [7], phenotyping the wild yeast *Lachancea quebecensis* [8], screening mutagenized strains of the unicellular alga *Chlamydomonas reinhardtii* [9], and generating *E. coli* synthetic genetic arrays [10]. The ROTOR HDA uses disposable plastic ‘repads’ of pins to print cells from one array to another in a single action. Long-pin repads can be used to transfer material to or from microtitre plates containing liquid cultures and to or from solid agar lawns in rectangular dishes (such as Singer’s own proprietory ‘PlusPlates^™^’). 96 spot arrays can easily be combined into 384 arrays, or 384 into 1536, by superimposing each source array with an offset. Larger arrays can easily be broken down into multiple smaller arrays by reversing that process.

Microbial growth can be quantified in liquid media in microtitre plates, and analysed using packages such as GrowthRates [11] or GATHODE [12]. It can most accurately be measured by measuring turbidity using specialist machines such the Bioscreen C (Oy Growth Curves Ab Ltd) [15, 16] which holds two 100× 600 μl sample plates with a proprietary honeycomb format meant to create more even heat distribution through the samples. However, it is challenging to simultaneously inoculate these hexagonal-grid wells from the rectangular-grid high-density ROTOR arrays, and to find a level of agitation sufficient to prevent flocculation that does not introduce bubbles or risk cross-contamination between wells. Another solution, the oCelloScope (Philips), uses standard microtitre plates but can only read 96 wells at once. The Omnilog (Biolog) can handle 50x 96-well plates simultaneously, but at a considerable cost and footprint.

It is becoming more common and is certainly more convenient to measure the growth of whole colonies growing on solid agar lawns in rectangular dishes. This is usually achieved by feeding digital snapshots, taken intermittently, into sophisticated image-analysis software, such as the Screenmill Colony Measurement Engine [14] (as open-source macro for the Java-based ImageJ software), Colonyzer [15] (a command-line app written in Python 2.7), YeastXtract [16] or the Colony-Live imaging system [17]. Specialist instruments have now become commercially available which perform this imaging and analysis in an automated fashion, for example the Phenobooth (Singer Instruments, Somerset, UK). Such an instrument greatly simplifies the process, and incorporating incubation and imaging functions into the same device overcomes limitations in the number of timepoints at which measurements are taken. However, image-based solutions remain vulnerable to confusion by specks and bubbles, particularly when measuring small colonies.

To overcome problems with these existing approaches and save time and money, we developed an alternative technique which uses a commonplace microplate reader (BMG Labtech’s FLUOstar Omega, which can incubate between 25 and 45°C and accommodate 96, 384 and 1536 plates) to measure the thickness of colonies printed with the ROTOR at 20 minute intervals. We have demonstrated that this measure is a reliable proxy for overall colony size. We developed the PHENOS software, with a graphical user interface (GUI), to run on Windows PCs alongside the microplate reader’s own software. This software combines multiple output files from the microplate readers and automatically generates a variety of data visualizations, and output files containing various growth curve summary values. The software is written in Python 2.7 and so could be adapted to other operating systems without great difficulty. It was designed to make it relatively easy to add new parser classes to recognise and read new formats of input files that might be generated by different microplate readers. Alternatively, users may prefer to write their own code for analyzing the data, or may find their microplate reader’s native software sufficient for their purposes.

We validated the PHENOS pipeline by comparing its measurements to two additional low-throughput means of measuring colony population: use of a cell counting chamber under a microscope, and absorbance in liquid media using a spectrophotometer. Comparing both absorbance methods to the manual cell count, we found PHENOS to be of comparable accuracy to the spectrophotometer. We also used this data to empirically derive a formula to estimate cell count from PHENOS readings. Because PHENOS measures absorbance through the thickness of a colony (a roughly cylindrical volume) as a proxy for overall colony size (a roughly hemispherical volume), there is not a straightforward linear relationship between the absorbance readings and cell count.

For additional validation and to demonstrate one of its potential uses, we used PHENOS measurements to reproduce a Quantitative Trait Loci (QTL) analysis previously undertaken using conventional liquid media growth measurements. QTL analysis is a powerful approach to understanding complex genetic architecture by correlating phenotypes with genotypes [18]. It can be substantially automated through the use of large arrays of genetically diverse microorganisms.

With sufficient genetic markers determined (e.g. by sequencing) for each strain in the array, growth phenotypes and genotypes can be passed into the statistical analysis package R/qtl [19]. Previously our lab described a set of 576 genetically-diverse haploid F1 *S. cerevisiae* strains derived from crosses between four geographically-diverse parent strains, chosen because they encapsulate much of the genetic diversity discovered in a genomic survey of *S. cerevisiae* [20]. The parents were YPS128 (herein designated ‘A’ for American), DBVPG6765 (‘E’ for European), Y12 (‘S’ for Sake) and DBVPG6044 (‘W’ for West African), each made genetically tractable with knockouts of the *URA3* uracil-metabolism gene and *HO* gene (thus preventing mating type switching and keeping strains haploid) [21]. From each of the six pairwise crosses between these parental strains, 24 tetrads were dissected, yielding 96 haploid F1s. These progeny are genetically and phenotypically extremely diverse. They were genotyped at over 200 markers using high-resolution melt quantitative PCR [22] to distinguish parental allele types, and then phenotyped for up to 23 different stress conditions in liquid media. 82 significant QTLs were described in total, including 4 distinct regions for resistance to the herbicide paraquat [23]. We phenotyped the same panel on solid YPD media containing 500 μg/μl paraquat and used the PHENOS pipeline to automatically generate data for entry into R/qtl [25]. We found the PHENOS method reproduced the QTLs from the earlier study but showed greater sensitivity as 18 additional regions were identified.

We consider a couple of the distinctive characteristics of PHENOS growth curves: an initial drop in absorbance is commonplace, and as colonies grow and absorbance measurements increase, variance in those measurements also increases. We investigate these properties experimentally and report their probable causes in our discussion and conclusions. Neither affects the utility of PHENOS, which produces growth curves that are both highly reproducible and accurate enough for statistically sensitive analyses such as QTL analysis.

QTL analysis on genetically diverse populations, by revealing genetic causes of phenotypic variation, is a valuable tool for determining the mode of action of drugs and stresses. But PHENOS will also be useful for measuring population trait variability in general, synthetic genetic interactions, assessing the quality of printed arrays, or for any of the other whole colony array techniques mentioned above.

## Methods

We will first describe, in general terms, the methods used in the PHENOS pipeline itself, including the punch-in technique for normalizing the amount of cells printed to each spot. For much greater detail, users can consult the guide which is available with the source code for the PHENOS software at https://github.com/gact/phenos. As part of this walkthrough, we also show examples of the visualizations that the PHENOS software produces. We then describe two methods used to validate this approach to phenotyping. One of these, using QTL analysis, provides a demonstration of how the PHENOS pipeline can be used in practice. We also describe a simple experiment to investigate properties of the PHENOS growth curves. Full details of all *S. cerevisiae* F1 segregants used are available at https://www2.le.ac.uk/colleges/medbiopsych/research/gact/resources/strain-resource.

### Overview

An overview of the experimental side of the PHENOS pipeline is given in Figure 1. The experimental plate is put into the microplate reader before any cells have been printed to it, in order to measure and later subtract out absorbance due to agar. As part of the printing process, starting cell masses can be partially normalized using the ‘punch-in’ technique described below. The microplate reader is set up to store each set of continuous readings as a different text file, and it is possible to generate growth curves even from intermittent readings. As summarised in Figure 2, the PHENOS software then gathers information from the user about the experiment (for instance the treatment and the name of the array being used, which will be used to look up information about individual strains in the layout, which is stored in a separate file), combines multiple files taken from a single experimental plate, and provides visualizations and analyses. The PHENOS software is written for Windows (so that it can operate alongside the microplate reader software) in Python 2.7, and has a simple GUI-based interface which can be launched by double-clicking on a desktop icon. It stores readings and other experimental data in HDF5 databases to allow complex queries and comparisons between multiple experiments, for example to measure final growth levels as a treatment/control ratio for use as a phenotype in QTL analysis.

**Figure 1.**
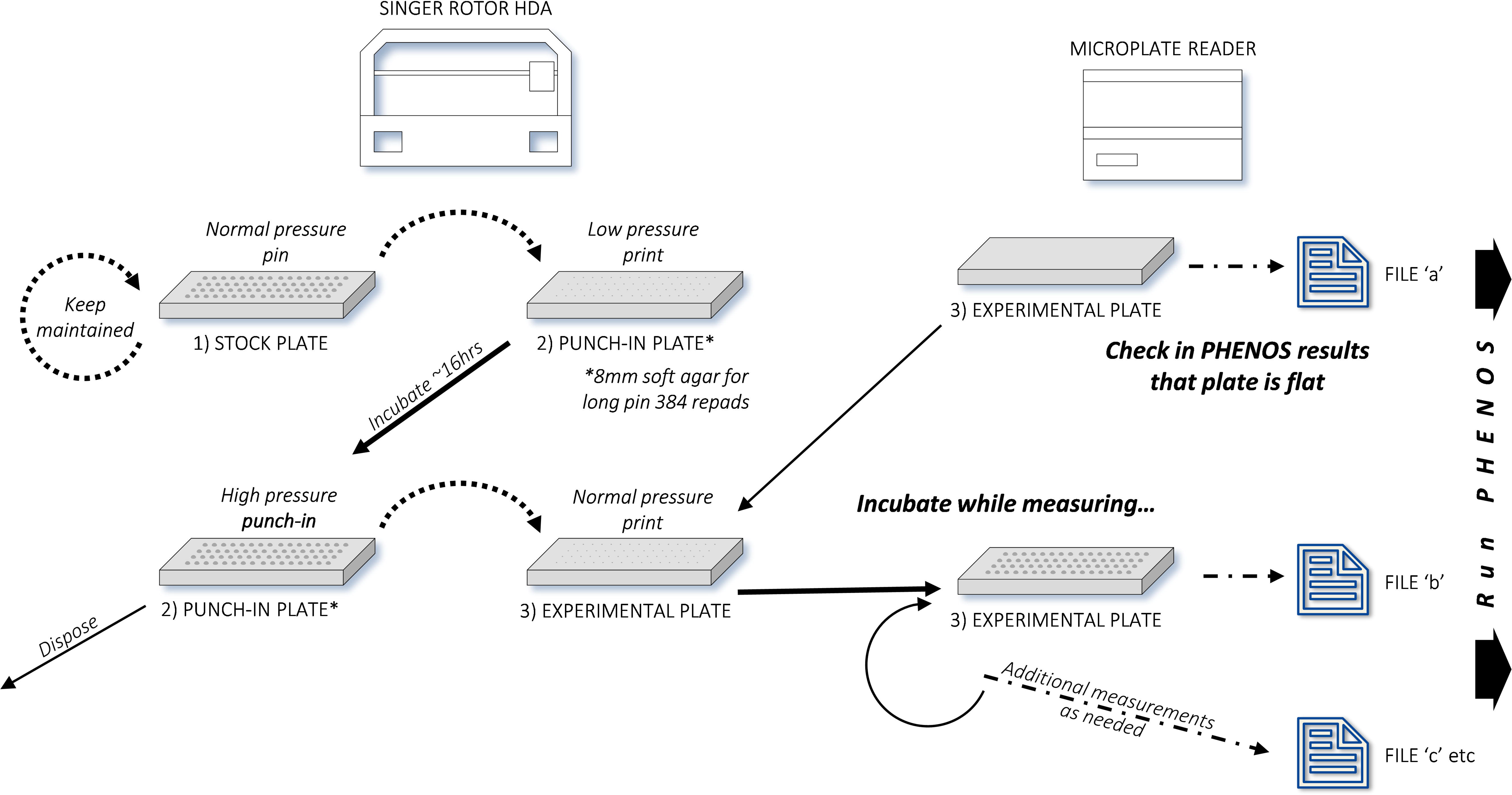
Overview of the PHENOS experimental workflow. We distinguish between three types of plates: stock plates for array maintenance, punch-in plates for normalization, and experimental plates for measurement.

**Figure 2.**
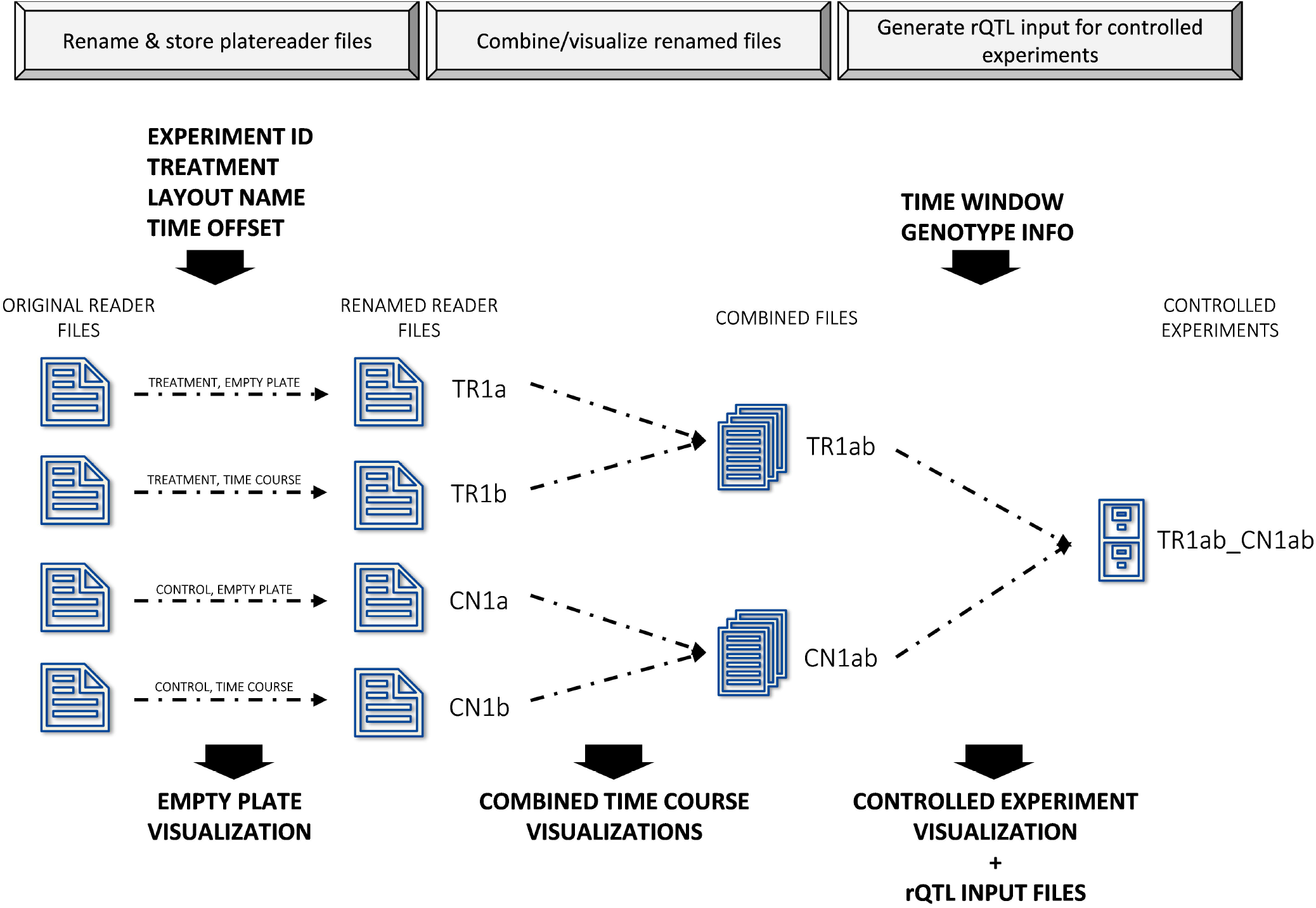
Overview of the PHENOS analysis workflow. The PHENOS software combines files and visualizes and compares experiments.

For our experiments, we typically compare normal growth (usually on standard rich YPD media) to growth under a phenotype-discriminating treatment (e.g. the same media containing a genotoxin). Our control experiments are normally arrayed at 384 density, with 2-4 replicates of each strain, on 40ml of YPD agar media: 10g/l Yeast Extract, 20g/l Bacto Peptone, 20g/l D-Glucose, 1% adenine solution (0.5% adenine in 0.05M HCI), adjusted to pH 6.3 using 1M HCI, then autoclaved with 20g/l Bacto agar. Treatment experiments are grown on the same media but with a treatment added (e.g. a toxic compound at a discriminating concentration, or a different incubation temperature). Absorbance readings (optical density, or OD) are taken at 600nm (on a FLUOstar Optima microplate reader, a 595nm filter can be used), every 20 minutes. Control experiments are run for approximately 65 hours (over a weekend) so they can be compared to as many treatment experiments as possible, as some treatment conditions do not produce discriminating growth phenotypes until much later. For many growth phenotypes, however, an overnight program run for ~16 hours is sufficient to discriminate between phenotypes based on final growth levels. Measuring full growth curves gives experimenters the clearest overview from which to judge the best summary values and time window to discriminate between different growth phenotypes. Once this judgement has been made, it is possible to increase productivity by taking only intermittent ‘snapshot’ measurements of each plate at the appropriate points. Our snapshot protocols take readings of a plate four times in rapid succession (taking under 20 minutes for a 384 array), in order to average over the measurement variance observed at high absorbance values, a phenomenon discussed in more detail below.

Examples of the visualizations produced by the PHENOS software are given in Figure 3. Additionally, the PHENOS software includes the option to generate time-lapse mp4 movies of colony thickness measurements, which are an effective means for calling attention to any location-related biases in growth. For each experiment (combining any number of sets of measurements of one experimental plate), PHENOS also generates a text file summary of the combined experimental data, a text file containing summary values of the curve for each colony and, if genotype data has been provided, a text file containing both summary values and genotype data suitable for analysis using the R/qtl package.

**Figure 3.**
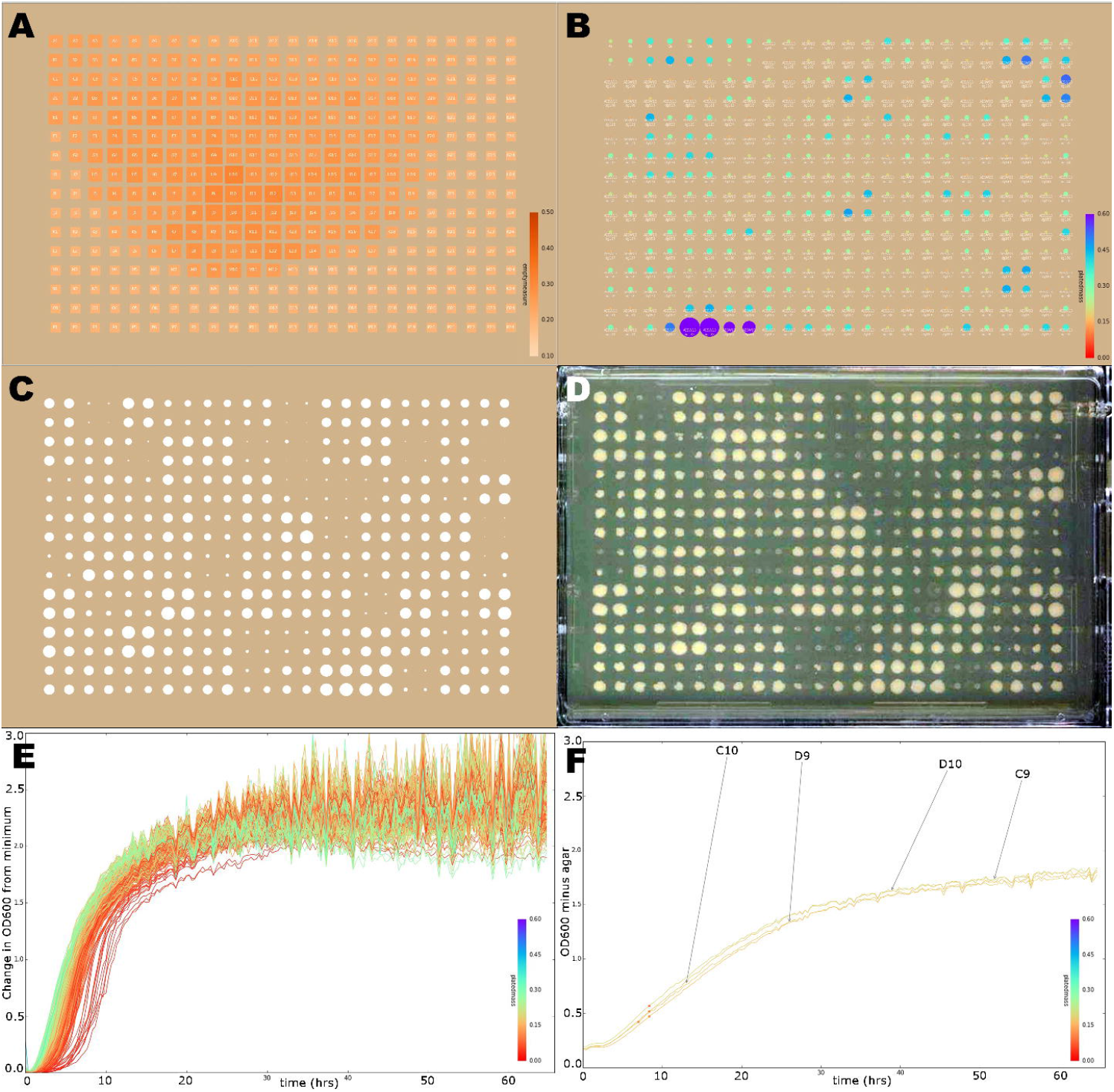
Examples of visualizations produced by the PHENOS software. The colour bars and some of the labels have been enlarged for this publication. **A)** Agar thickness of an unprinted plate, showing a low-quality plate with a bulge due to inadequately mixed media. For a good quality plate, all squares would be the same colour (the colour scale shows absorbance at each location in the array—darker colours signifying thicker agar—and is comparable between different visualizations). There should be no pattern to the size of squares either: the largest and smallest squares are the highest and lowest readings on that plate respectively. Squares are also labelled with the array position, although those labels are not legible in these small reproductions. **B)** Printing quality, based on initial readings immediately after printing, with agar subtracted. As with A, the colour scale is absolute and comparable between different experiments, but the spot scaling shows where the largest and smallest printed masses on the plate are located. Each spot is labelled with the strain name, which is legible on full size images. **C)** Final measurements of a plate as measured by the microplate reader (radius being proportional to OD_600_ with agar absorbance subtracted), next to **D)** an actual image of the same plate. In this case, there are four replicates of each strain. **E)** Control growth curves for all 384 colonies on a YPD plate, coloured by the initial printed mass as described for B. In a successful treatment experiment such as is shown in C & D, there would be much greater diversity in these curves. **F)** Replicate plots, showing growth curves of all four replicates of a single strain for which growth has been equally suppressed (compare to E) by hydroxyurea. Dots on these curves indicate where the PHENOS software has calculated the inflection point of each curve to lie, i.e. the point of maximum growth. Curves are also labelled with the array position of the colony.

The PHENOS software is written with a flexible architecture that allows users to configure which visualizations and summary values are included in the output by editing a text configuration file called ‘config.tx’. Each summary value is handled by a specific ‘phenotype calculator’ subroutine, and new ones can be written into the software with relative ease. These calculator routines are named in ‘camel case’, following programming convention. The currently available calculators are shown in Figure 4 and listed in ‘config.tx’. The calculators used by default can be changed by editing this file, or particular calculators can be specified for particular treatments. The PHENOS software can be used to pair a treatment experiment with a control experiment, creating a ‘controlled experiment’ database entry that allows for more complex phenotype calculators that compare the two curves (e.g. ‘TreatmentRatioCalc’, which is the most commonly used for our QTL analysis).

**Figure 4.**
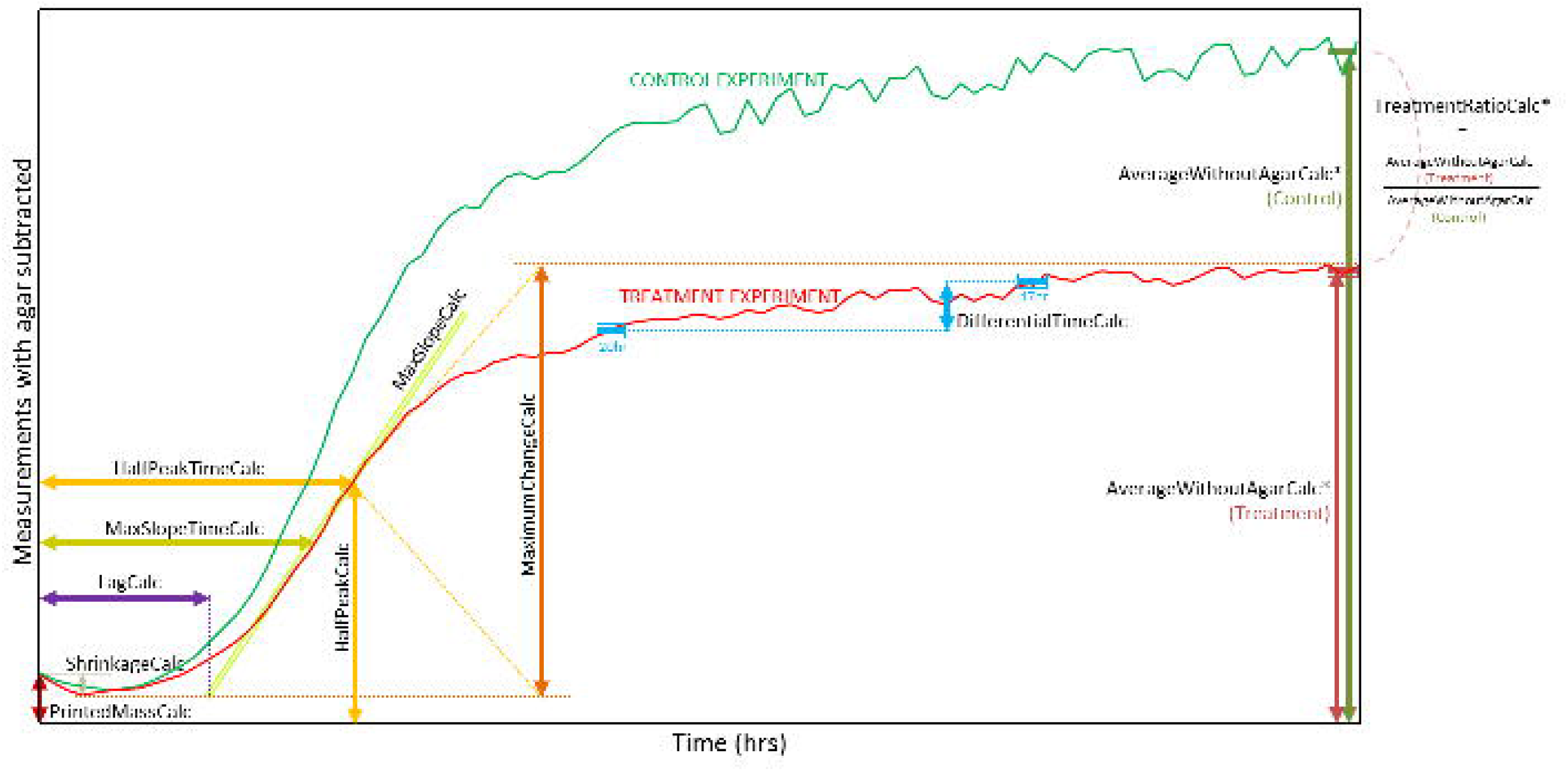
‘Phenotype calculators’ within the PHENOS software. Each generates a different summary value. Some, indicated with asterisks, e.g. AverageWithoutAgarCalc, require a particular time window to be specified by the user: for a given curve, all timepoints within that window are then averaged, to account for the variance usually observed at higher absorbance readings. Distinctive properties of PHENOS curves, such as initial reductions in absorbance, are discussed within the text and Figure 6.

### Normalization by the punch-in technique

For best results, there should be as little variation as possible in the amount of cells printed to each position in the plate to be phenotyped. We have developed a ‘punch-in’ technique to normalize these printed masses, which is summarised in Figure 5.

**Figure 5.**
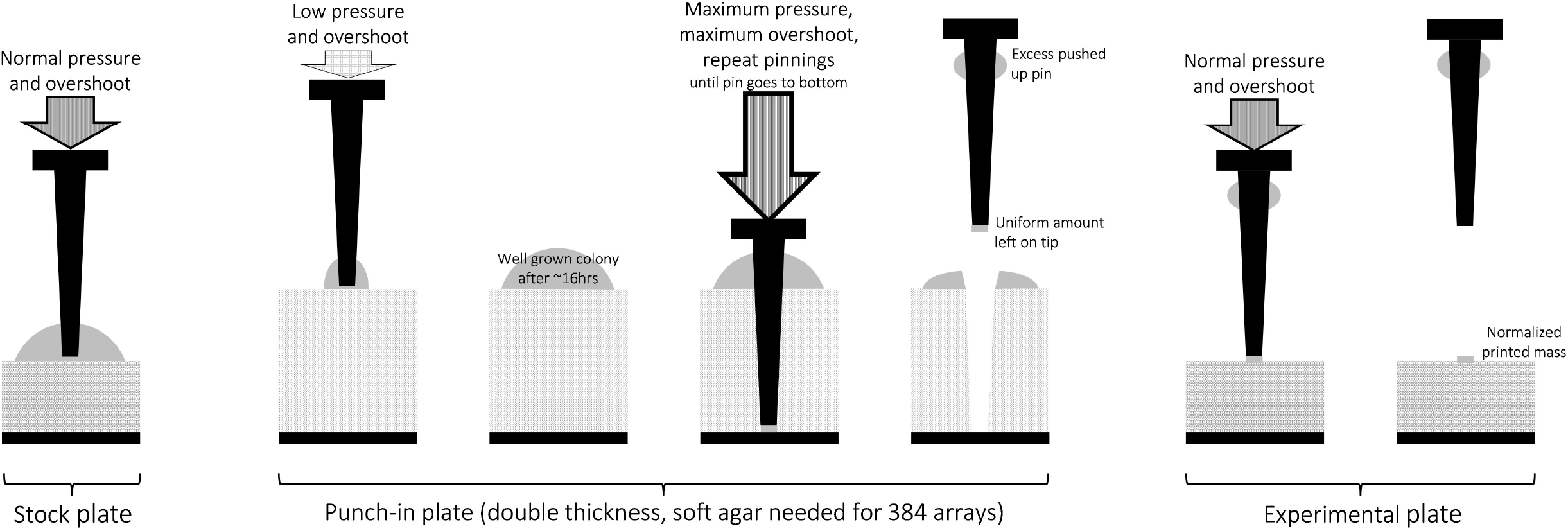
Punch-in technique for normalizing printed cell masses.

We create special single-use ‘punch-in’ plates with double the usual thickness of agar medium (8mm rather than 4mm), and allow the colonies to grow on it overnight. We use long-pin repads and turn the ROTOR pressure and overshoot settings to maximum, in order to drive the pins into the agar and push excess cell mass up the sides of the pin, leaving a more uniform amount on the tip, which is then printed to the target (experimental) plate as normal. When working with 384 arrays and repads, the ROTOR cannot deliver sufficient pressure to punch a 384 repad into normal agar. Therefore, a soft agar punch-in plate, with 25% of the usual agar, must be prepared (and used within 24 hours). The yeast array must be printed to this plate, using low pressure and overshoot settings so as to not punch in prematurely. Only 384 repads should be used to print to soft agar plates because 96 repads will punch in to them immediately even at the lowest possible settings, which will prevent proper punch-in normalization when they are later used as a source plate. Printed punch-in plates should be incubated for ~16 hours before use, and then used as a source plate almost immediately. Pinning settings for this source plate must be set to maximum pressure and overshoot, and sufficient repeat pinnings to drive the pins all the way to the bottom of the soft agar precisely once. The detailed PHENOS guide provides extra tips for mastering this technique, and it should be practised before being used for actual experiments. We have observed differences in normalization efficiency are most often due to different makes of agar or inconsistent media pH producing variation in agar hardness, or due to punch-in plates not being thick enough (we recommend 8mm).

### Cell count validation

We created multiple 96-density arrays on solid media of a single haploid strain with weak flocculation (EW01fc403), and induced a range of colony sizes by incubating the plates from one edge and for different lengths of time. Each plate was measured using PHENOS, with a ‘snapshot’ program to take four successive readings, and immediately afterwards colonies of different sizes were extracted along with a plug of agar by using a 1 ml pipette tip with 15-20 mm cut off. Colony and agar were deposited into 1 ml deflocculation buffer (50 mM KAc, 5 mM EDTA buffer, pH 4.5), and cells rinsed from the agar by thorough pipetting. The solution was then removed to a cuvette, leaving agar behind, and blanked readings taken in a Helios Gamma spectrophotometer (Thermo Scientific). Thereafter, 5 μl was transferred to a Neubauer improved cell counting chamber for counting at 40x magnification. Number of cells = 10000 × C × (25/S) where C=cells counted, S=subsquares considered. In some cases, freshly printed colonies were extracted into only 100 μl deflocculation buffer for cell counting without also being read in the spectrophotometer. Data was tabulated in Microsoft Excel, and microplate reader measurements were adjusted to remove absorbance due to agar, and to average readings from the snapshot program. For spectrophotometer readings with OD_600_>1, 1:10 dilutions were used instead and results multiplied by 10 to account for known inaccuracies at high optical densities.

### QTL validation

The 576 F1 *S. cerevisiae* strains previously described [23], comprising six crosses which we redesignated as ‘AE’, ‘AS’, ‘AW’, ‘ES’, ‘EW’ and ‘SW’, were re-arrayed as three 384-density arrays, each containing two of the crosses, with two biological replicates of each. All were incubated at 30°C for at least 16 hours, both with and without 500 μg/μl paraquat mixed into the YPD agar media (after autoclaving), and measured every 20 minutes. For each replicate, we determined the average ODeoo (with agar absorbance subtracted) at ~16 hours as a fraction of the control growth (without paraquat) using the TreatmentRatio phenotype calculator in the PHENOS software. Biological replicates were kept separate rather than averaged together in order to improve statistical power. Taking these biological replicates as input, R/qtl was used to calculate genotype probabilities at intervals of 1 centiMorgan, perform single-QTL analysis on the input data, run single-QTL analyses on 1000 permuted datasets, and estimate a LOD threshold at 5% significance level based on those permutation analyses. Before being input to R/qtl, each permuted dataset was generated by stratified permutation within tetrads, ensuring that no sample was permuted with its biological replicate. QTL peaks were identified at loci with LOD values exceeding the threshold, and 1.5-LOD support intervals were estimated by extension from the QTL peaks in either direction until the LOD value fell by more than 1.5 LOD units below the peak LOD value.

### Analysis of measurement variance

In order to assess the impact of material uniformity on variance in successive measurements, we took multiple successive readings of a range of organic and non-organic substances with different textures and levels of uniformity or granularity: yeast colonies now in stationary phase, artificial spots made from woodglue and yeast extract mixed with biro ink, and layers of printer paper, opaque plastic sheeting (cut from disposable weighing boats), and low-density 2mm foam packing material. The standard deviation of multiple successive measurements was calculated for each material type.

## Results

### Early dips in PHENOS growth curves

A common peculiarity of PHENOS curves is a slight dip that is often observed over the first hour of incubation (clearly visible in Figure 4). We attribute this to microscopic bubbles which we have observed in freshly printed spots (Figure 6), which behave like lenses and diffract the microplate reader’s light source until they have evaporated away. While curves can be normalized against each other according to their minimum values, we observe in replicate plots that curves are usually well aligned over most of the time course if the shrinkage is simply ignored.

**Figure 6.**
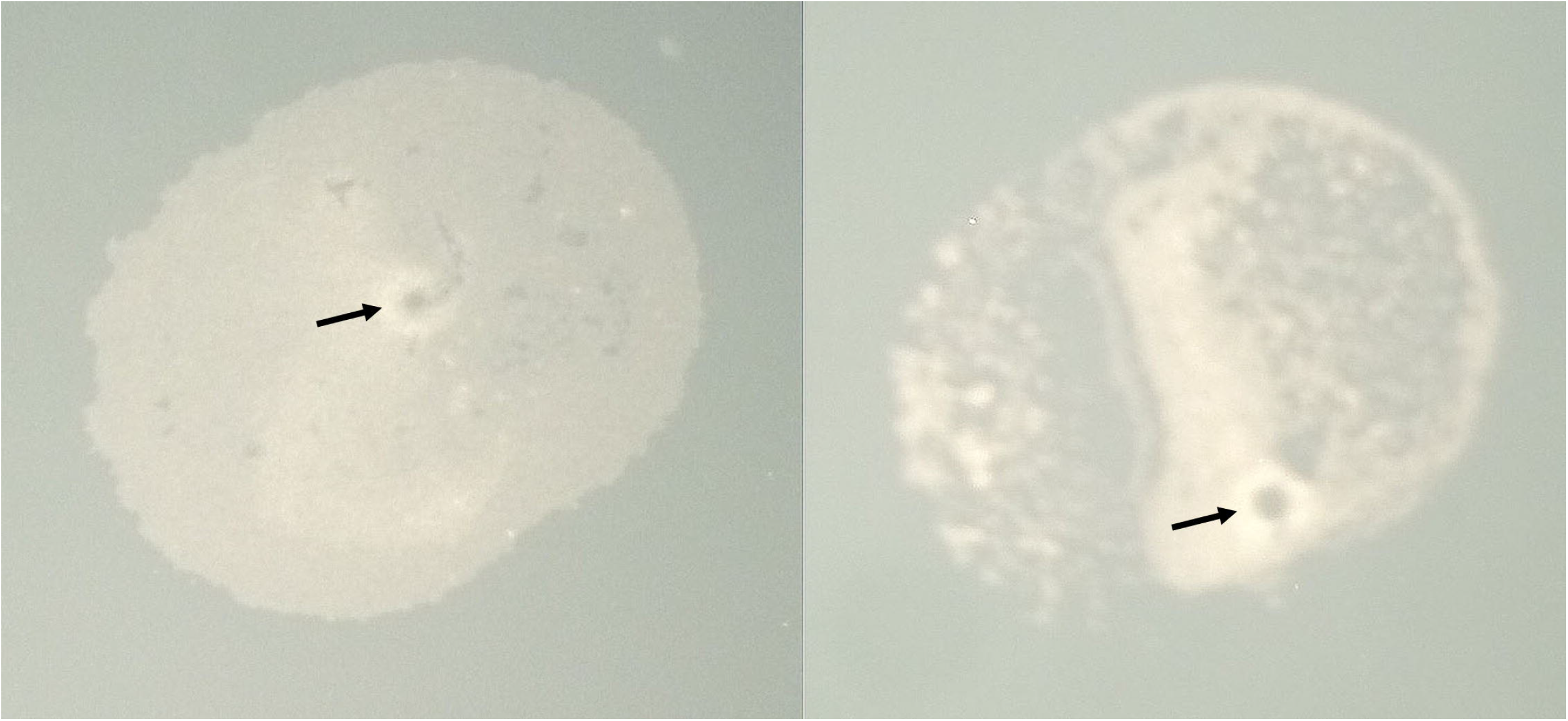
Bubbles in printed cell patches. These evaporate over the first hour after printing, leading to initial dips in PHENOS growth curves.

### Cell count validation

A range of differently sized colonies were measured in multiple ways, as described in the corresponding methods section. The measurements and calculations are in Additional File 1, and are displayed in Figure 7. To fit curves to each set of comparisons – spectrophotometer vs cell count, and PHENOS vs log_10_ (cell count) – Excel’s LINEST function was used to determine parameters that minimize the sum of least squares, giving best curve fits relating A (average measurement with agar subtracted) to P (predicted cell count). With parameters rounded, the best fit for PHENOS is P=77900e^(1.76A)^, while for liquid spectrophotometry the relationship is P=8000(725A+7). For average predicted cell counts below 6 million, corresponding to a microplate reader measurement (without agar) of ~2.5, the predicted cell counts for the two methods show linear agreement with R^2^=0.77. Following the statistical approach of Bland & Altman for comparing two methods of measurement [24], it can be seen in Figure 8 that at higher absorbance measurements both methods show comparable degrees of inaccuracy, although the PHENOS formula tends to underestimate the size of large colonies.

**Figure 7.**
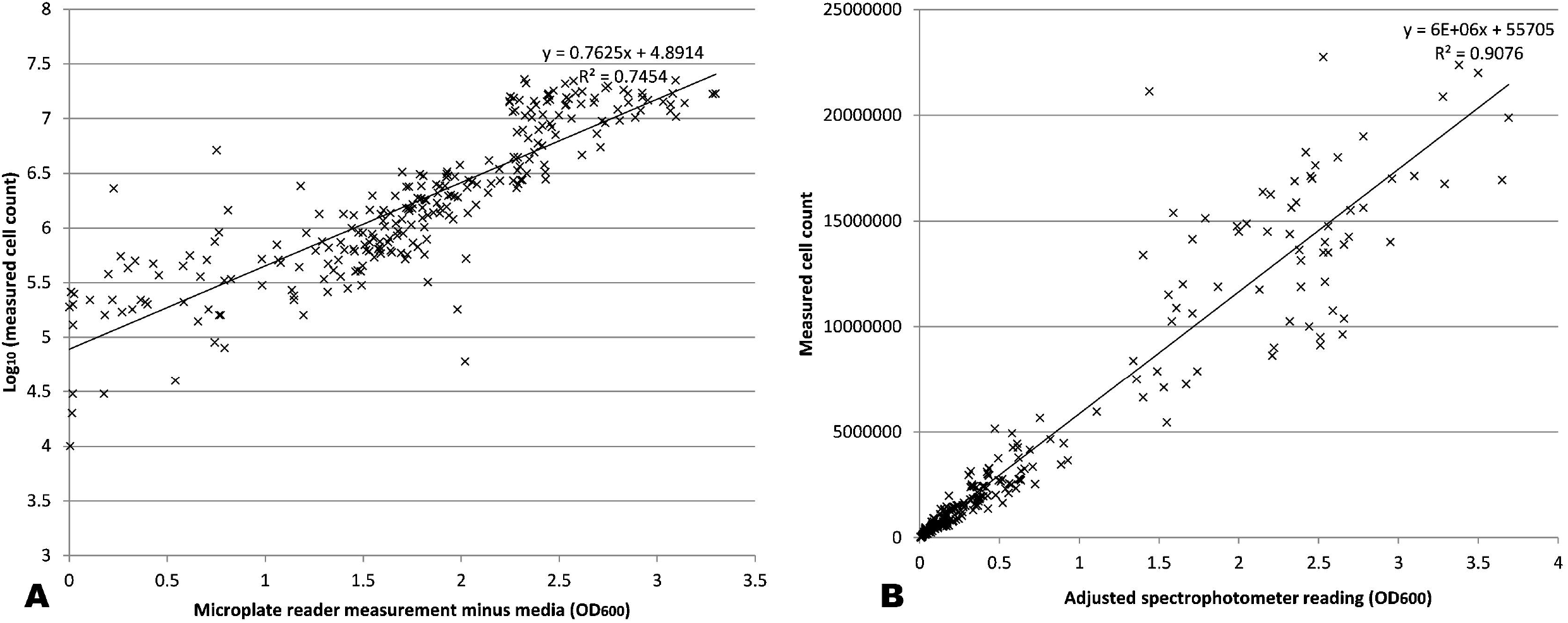
Accuracy of PHENOS colony measurements. **A)** Microplate reader measurements plotted against cell counts. **B)** Liquid spectrophotometry measurements plotted against cell counts; for initial readings >1, 1:10 dilutions were read and their results multiplied by 10 to empirically adjust for non-linearity at high absorbances. Together, these measurements were used to empirically determine formulae for predicting cell counts from measured values.

**Figure 8.**
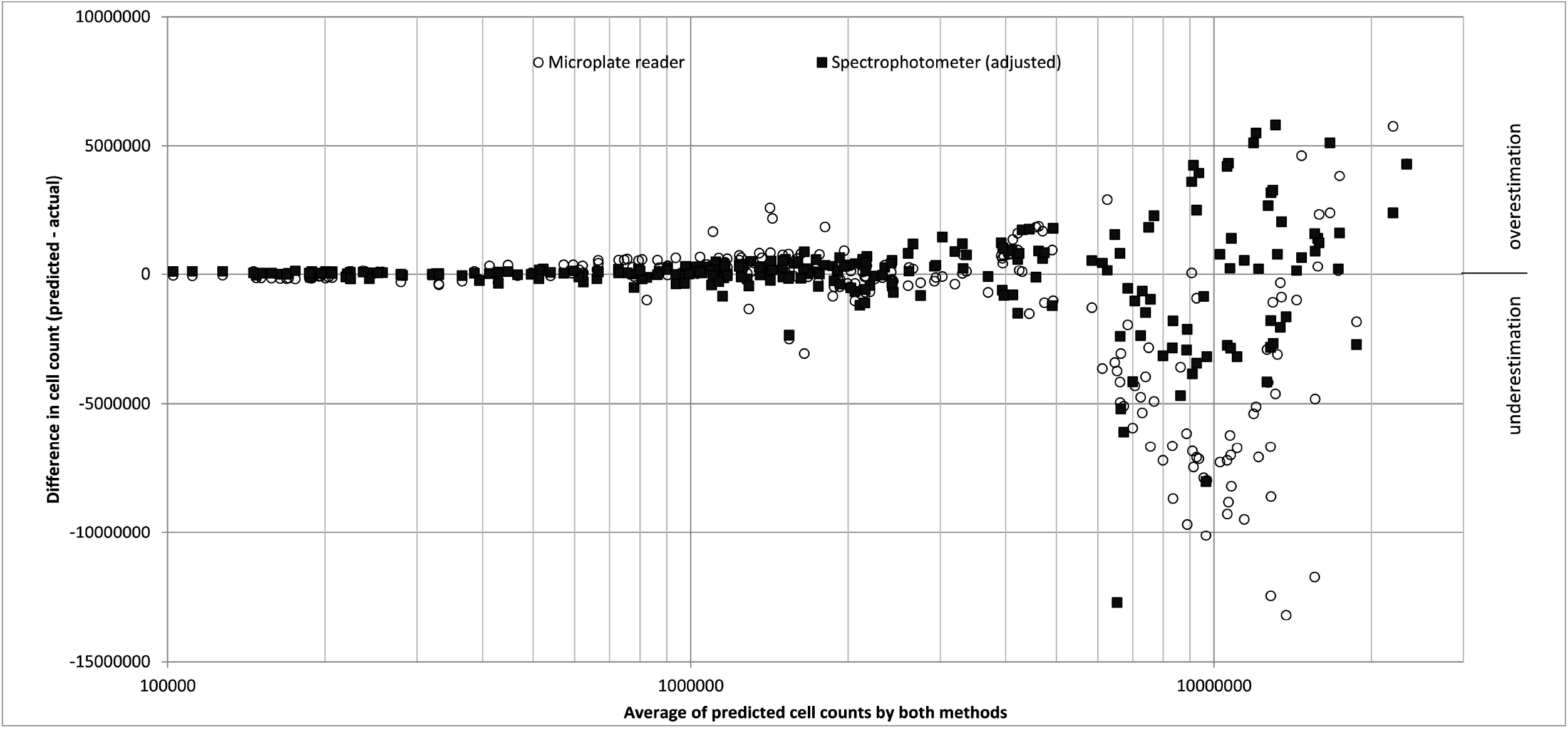
Comparison of predicted cell counts by two methods: microplate absorbance and conventional liquid spectrophotometry, both at OD_600_.

### Variance in measurements at high absorbance values

We observe that higher absorbance readings, above 2.0 (typically reached in 10–20 hours on YPD at 30°C) exhibit greater variation between successive measurements, and often in a manner that is synchronised across most colonies on a plate (as can be seen in Figure 3E), but with no discernible periodicity. This variance is equally noticeable with more frequent measurements, with and without incubation, and whether or not the reader is covered up to limit any light leakage or heat flow. We tested the hypothesis that this might be due to microscopic differences in the starting position of each pass of readings, and therefore would be more pronounced for colonies with more elaborate morphologies or, in general, with the granularity of the material being measured. We took multiple successive measurements of a range of organic and non-organic substances with different granularity, but observed no difference between them (Figure 9, Additional File 2).

**Figure 9.**
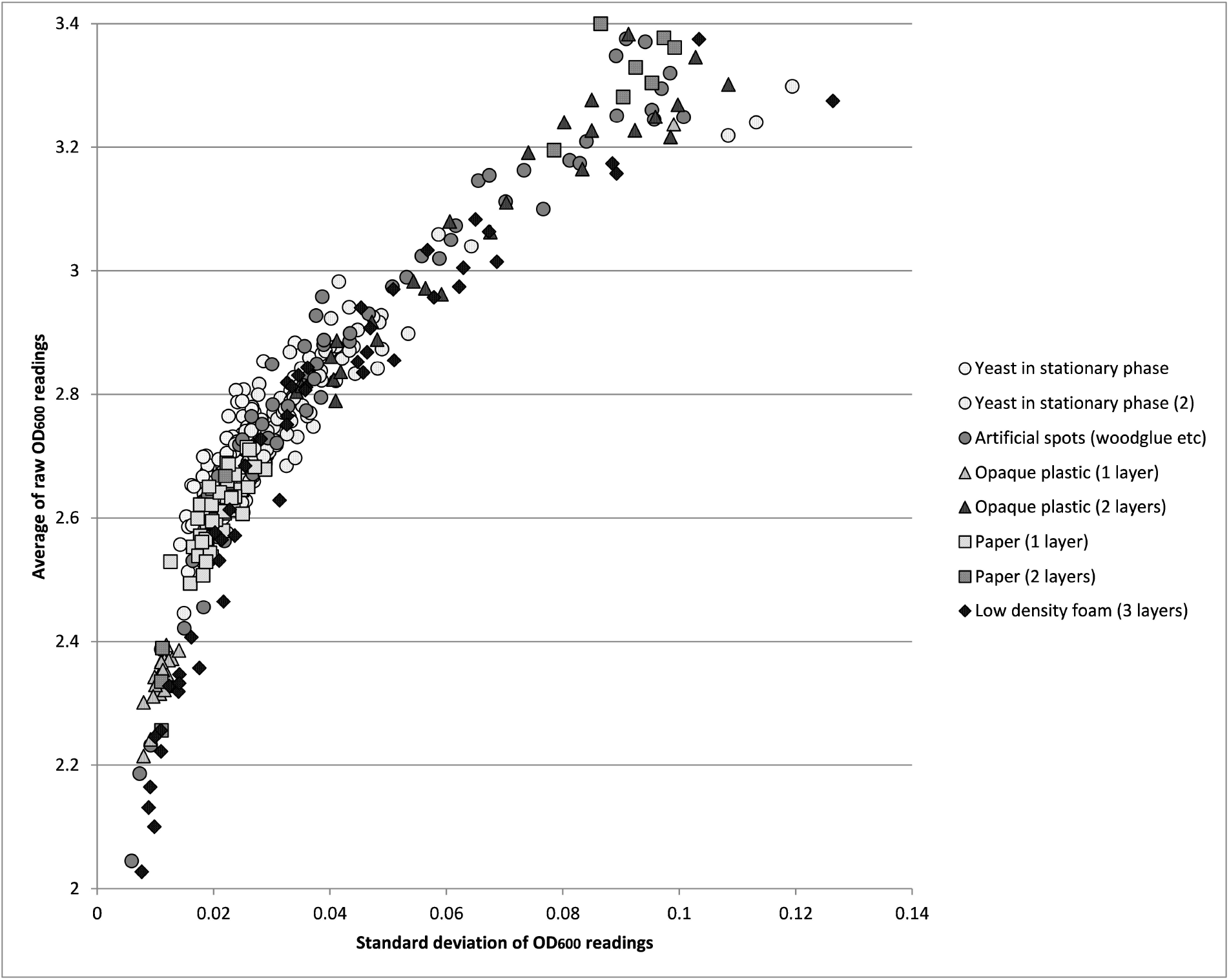
Variation in multiple successive readings of a variety of materials. This variation is a function of overall average absorbance, but barely affected by different degrees of granularity.

These results indicate that reading variation at high absorbance values is not attributable to positioning error and granularity. We conclude that this is simply an unavoidable hardware limitation, though easily overcome by averaging readings within a time window, as in our snapshot protocols.

### QTL validation

PHENOS was used to derive treatment ratios for the 576 F1 haploids growing with and without 500 μg/μl paraquat. Sample curves are shown in Figure 10.

**Figure 10.**
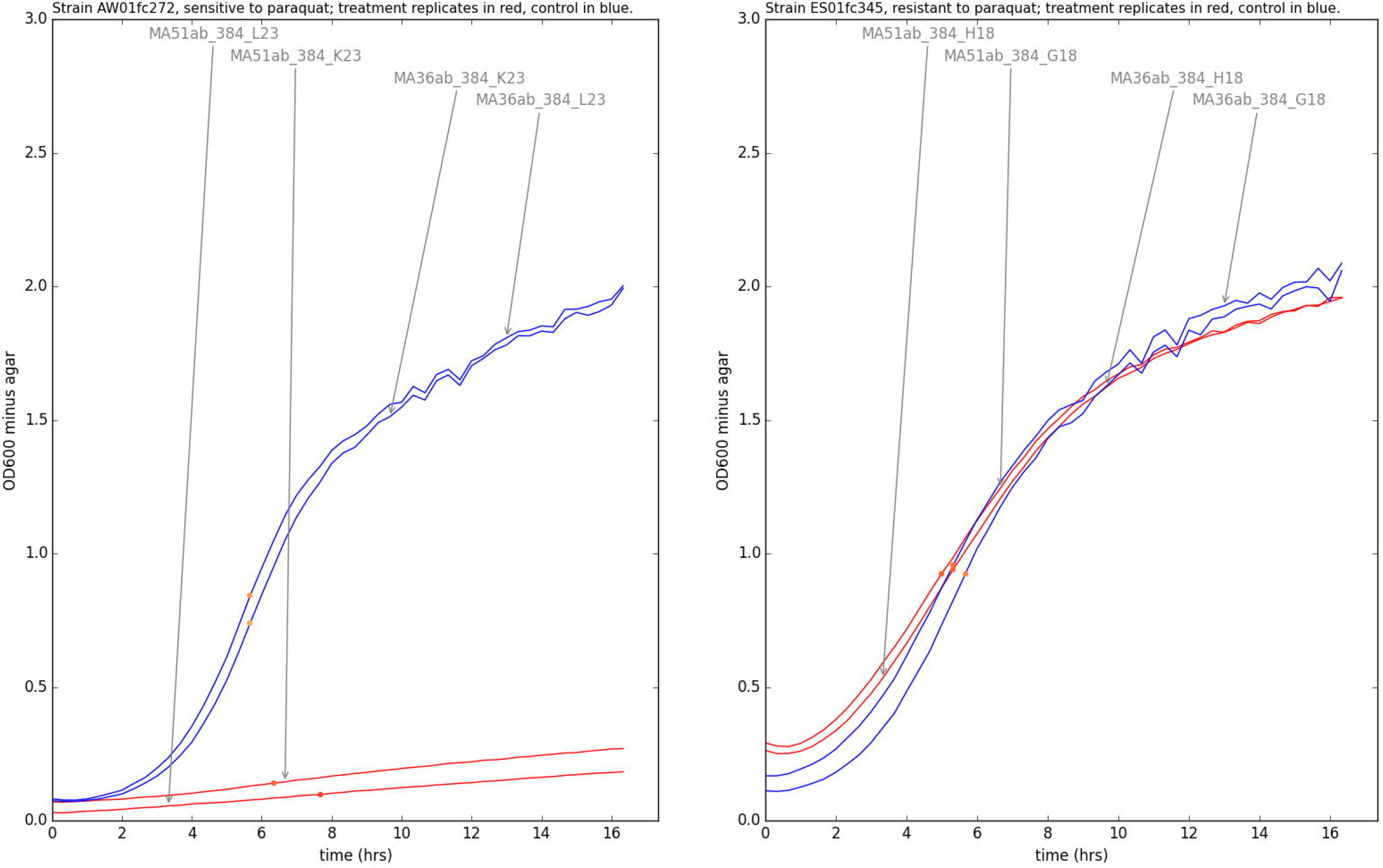
Sample curves from two strains showing extremely different growth responses. Growth on solid media containing 500 μg/μl paraquat (red curves) compared to normal YPD (blue curves). Curves are labelled with experiment IDs and plate positions, and dots indicate PHENOS-calculated inflection points.

QTL analysis with R/qtl yielded 61 QTLs overall, covering 22 distinct regions, including all four found in the previous study [23], with comparable LOD scores, as illustrated in Figure 11 (see Additional Files 3A and 3B for dataset and original measurements respectively). The original study in liquid media, and this reproduction of it, both had low resolution due to large haplotype blocks and low density genetic markers. Much greater resolution, down to individual SNPs, can be achieved by repeating a process of crossing and sporulation to create advanced intercross lines with a high level of genomic admixture, and generating much larger numbers of markers through whole genome sequencing [25].

**Figure 11.**
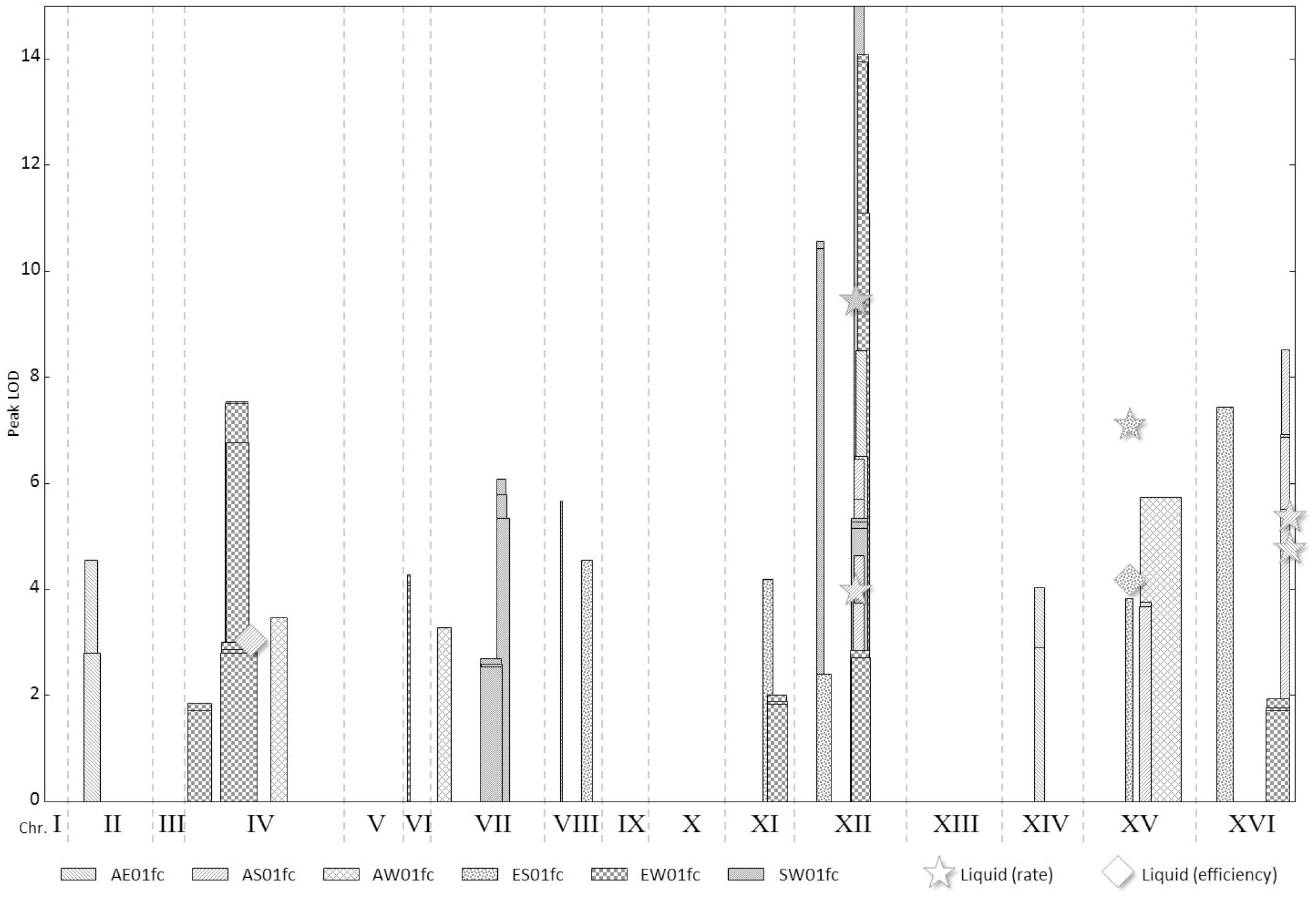
QTL validation. *S. cerevisiae* QTLs for paraquat resistance, determined using PHENOS, compared to QTLs found in the same set of strains grown in liquid media. In both experiments, 576 strains were phenotyped, each an F1 progeny from one of six different crosses of four geographically diverse parental strains (A= American, E= European, S= Saki, W=West African). Different crosses produce different QTLs, though one QTL on chromosome XII is common to all the crosses. The results show that PHENOS reproduces the liquid media QTLs and is more sensitive.

## Discussion

Despite the reading variation at higher absorbance values described above (and equally problematic in liquid spectrophotometry) PHENOS can provide accurate scores of colony size across a very wide range of sizes, including very small ones that might not be picked up at all by imaging-based approaches. Analyses of inter- and intra-plate variability (Figures 9 and 10 respectively) demonstrate that this is a robust and highly reproducible assay.

We have already described and investigated a couple of quirks of PHENOS growth curves. These make it difficult to reliably fit the curves to conventional logistic growth curve models [26]. Additionally, we have observed some non-standard yeast growth patterns in response to various genotoxins. Strong doses can produce curves that that show growth initially but then trend downwards as cell death and dispersal thins the colony. Some compounds with lower stability, such as methyl methanesulfonate, can produce curves which plateau and maybe fall before then entering a second phase of growth after the survivors begin to proliferate. These are interesting responses and worthy of study even though they cannot be parameterized as conventional logistic growth models. For the purposes of QTL analysis, the most powerful phenotype measurement is the summary value that shows the greatest variation between sensitive and resistant strains, and is least influenced by experimental factors such as normalization efficiency, or least prone to miscomputation. This may vary on a case by case basis, depending on the growth patterns concerned, so it is best to view the full growth curves before deciding on the best summary values (and time windows) to use.

As shown in Figure 4, the PHENOS software allows for a range of summary values to be generated from a PHENOS curve, including ones which are analogous to the efficiency (final growth), rate (maximum slope), and lag (growth delay) parameters commonly derived from logistic growth curves in liquid media, [26, 28]. The ‘MaxSlopeCalc’ value, analogous to growth rate, can be miscomputed when there are multiple growth phases, so replicate plots should be consulted to confirm that inflection points are being correctly identified. The ‘LagCalc’ value, analogous to growth lag, often correlates with the initial absorbance of cells right after printing (the ‘printed mass’). For this reason, by default, PHENOS generates a scatterplot visualization comparing the two values for each curve so that any correlation becomes evident. For the key purpose of discriminating between strains, the growth level in a hand-picked time window (‘AverageWithoutAgarCalc’), and its cross-experiment derivative (‘TreatmentRatioCalc’) are usually the most direct and reliably computed values.

So-called ‘edge effects’ are a common concern of working with microbial arrays. The colonies around the edge of an array might grow differently due to differences in heat-dispersal (in liquids) or nutrient availability (on solid media). In the solid media arrays used for PHENOS, we only observe significant edge effects after about 40 hours in arrays growing on normal YPD media. At this point peripheral colonies continue growing for longer due to greater availability of nutrients at the edge of the array, but we see no evidence of significant differences in earlier growth, or on plates where growth is limited by treatments instead of nutrient availability.

In some circumstances PHENOS may not be the most suitable strategy for growth phenotyping. If a treatment compound is expensive it will be inefficiently deployed by being dispersed in 40 ml of solid media. Reducing the amount of media to 20ml per plate may mitigate this, but can introduce artefacts due to the agar slightly contracting from the edges of the plate during incubation. Alternatively, it might be possible to dispense hot agar into the wells of a microtitre plate but care would need to be taken to avoid bubbles and unevenness (possibly with the use of additives such as surfactants) that would compromise printing quality, and this would be harder in 384-well plates than in 96.

Any treatment compound must be soluble of course, and sufficiently stable in solid media, and if it affects the pH too strongly it will also make the agar too soft. If the treatment compound affects the agar colour too much then it may increase the base agar absorbance readings so much that growth curves appear to plateau prematurely, because the maximum possible overall reading, for FLUOstar microplate readers at least, is 3.5. A typical *S. cerevisiae* colony at maximum size in a 384-array growing on normal YPD agar will register an OD_600_ without agar of up to 2.5, so the agar absorbance must be less than 1 when working with this organism. Complex phenotypes such as colony morphology characteristics cannot easily be measured by PHENOS in its current state. Furthermore, we expect that growth on solid media will not always reflect growth under liquid conditions. The latter may be more relevant to applications in brewing and wine-making, although solid media growth is likely to be much more similar to how yeast grow in the wild.

We have found 384 arrays strike the best balance between scale and reliability but the ROTOR is capable of printing 1536 arrays and, with a minor vendor adjustment, the FLUOstar Omega is capable of reading them. 1536 arrays are challenging to duplicate, especially if they contain diverse growth phenotypes, as slow-growing colonies are easily lost, often due to larger surrounding colonies keeping a repad pin from making full contact. Only short-pin repads are available for 1536 arrays, and their pins have a very small cross section which necessitates a much finer alignment between printing and reading coordinates than the machines can usually achieve. This problem may be overcome by using 4 long-pin 384 repads, each normalized from a different punch-in plate, but this reduces the convenience and speed of the method. Furthermore, the greater proximity of colonies in 1536 arrays substantially reduces growth speed and peak colony size, and brings forward the time point at which edge effects become prominent. Also, mating pheromone inhibition or similar effects may complicate growth in mixed arrays.

## Conclusions

In successfully normalized experiments, PHENOS produces proxy growth curves with a high degree of reproducibility between replicates (Figures 3F, 10), and with greater ease and automation than conventional methods. We have reported results that use *S. cerevisiae*, but within our lab the exact same approach has also been used to measure the growth of other yeast strains such as *S. eubayanus* CBS 12357^T^ [27] and *S. uvarum* CBS 7001 [28]. Nor need PHENOS be limited to working with yeast. We have briefly experimented with using PHENOS to measure *E. coli* growth (Thermofisher MAX Efficiency^™^ DH5α^™^ Competent Cells) on Luria Broth agar, and obtained growth curves with a maximum change in OD_600_ of 0.62 after 16 hours (Additional File 4). While less optically absorbent than yeast, this should still be perfectly sufficient to discriminate growth phenotypes. Provided that an organism can grow on a solid medium in the first place, PHENOS may prove a viable approach to growth phenotyping.

We have also briefly experimented with measuring whole-colony fluorescence in tandem with optical absorbance and have encouraging preliminary results, although extra software will be needed to handle the data files produced by this procedure.

For laboratories in possession of a colony arraying robot, colony thickness as measured by an incubating microplate reader is a very convenient and sensitive alternative to more expensive specialist solutions for quantifying growth. It sidesteps problems with imaging-based solutions such as poor contrast or bubbles or specks which can confuse image analysis, as well as the general difficulty of setting up a high-quality imaging and image analysis pipeline in the first place. It avoids problems with liquid phenotyping such as format conversion, unequal heat distribution, evaporation, and the technical challenge of fine-tuning agitation levels to prevent flocculation without introducing bubbles or cross-well contamination. Furthermore, there are clear safety benefits to having toxic treatment compounds contained within solid agar, and the same will be true if working with more dangerous organisms than yeast

## List of Abbreviations

OD: : optical density;
PHENOS: : phenotyping on solid media;
QTL: : quantitative trait loci;
SGA: : synthetic genetic array;
YPD: : yeast extract peptone dextrose growth media.

## Additional Files

**Additional File** 1 Validation by cell counting (xlsx). Raw data and calculations.

**Additional File 2** Testing with multiple materials (xlsx). Raw data and calculations.

**Additional File 3A** Validation by paraquat QTLs (xlsx). Full list of QTL intervals and their details.

**Additional File 3B** Validation by paraquat QTLs measurements (zip). Initial DAT and csv files.

**Additional File 4E** coli results (xlsx). Raw data and calculations.

## Declarations

### Ethics approval and consent to participate

Not applicable.

### Consent for publication

Not applicable.

### Availability of data and materials

The datasets supporting the conclusions of this article are included within the article (and its additional files). The PHENOS software and test files are available from https://github.com/gact/phenos/. under an MIT license. This software is written for Windows, and requires Python 2.7, along with the modules “scipy”, “numpy”, “tables”, “matplotlib”, “xlrd”, “brewer2mpl” and their dependencies. The yeast strains used in QTL validation are available on request.

### Competing interests

The authors declare that they have no competing interests.

### Funding

The work was funded by the University of Leicester under a Burgess Centre grant, with additional funding for consumables, genotyping and equipment provided by BBSRC grant BB/L022508/1 to EJL and by Wellcome Leicester ISSF award 097828/Z/11/B and Hope Against Cancer Small Research Grant to SSF. DG was funded by the University of Leicester under a Pritchard PhD studentship award. The funding bodies had no role in the design, interpretation, analysis, discussion or writing of the work.

### Author’s contributions

DBHB invented the technique, wrote the PHENOS software and drafted the manuscript. ND tested early versions and provided feedback. MA performed the paraquat phenotyping. TW provided help with R/qtl and Github. DG tested the software in a variety of contexts and drove improvements. EJL and SSF helped interpret results, assess outputs, and helped draft the manuscript. All authors read and approved the final manuscript.

## Acknowledgements

We wish to thank Maria Viskaduraki for statistical advice, and Tuğba Şan and Noor Bafana for testing aspects of the pipeline in its early stages.

